# Who’s who? Detecting and resolving sample anomalies in human DNA sequencing studies with *peddy*

**DOI:** 10.1101/074385

**Authors:** Brent S. Pedersen, Aaron R. Quinlan

## Abstract

The potential for genetic discovery in human DNA sequencing studies is greatly diminished if DNA samples from the cohort are mislabelled, swapped, contaminated, or include unintended individuals. Unfortunately, the potential for such errors is significant since DNA samples are often manipulated by several protocols, labs or scientists in the process of sequencing. We have developed *peddy* to identify and facilitate the remediation of such errors via interactive visualizations and reports comparing the stated sex, relatedness, and ancestry to what is inferred from each individual’s genotypes. *Peddy* predicts a sample’s ancestry using a machine learning model trained on individuals of diverse ancestries from the 1000 Genomes Project reference panel. *Peddy*’s speed, text reports and web interface facilitate both automated and visual detection of sample swaps, poor sequencing quality and other indicators of sample problems that, were they left undetected, would inhibit discovery.

**Software Availability:** https://github.com/brentp/peddy

**Demonstration (Chrome suggested):** http://home.chpc.utah.edu/∼u6000771//plots/ceph1463.html

## Introduction

Human DNA sequencing studies frequently involve the handling of DNA samples and associated manifests by multiple laboratories and individuals. Both whole-exome (WES) and whole-genome (WGS) sequencing protocols involve multiple DNA manipulations prior to sequencing. Each new procedure and handling is another opportunity for sample mix-ups, contamination, or mislabelling. Even a single DNA mix-up has the potential to destroy discovery and diagnostic power. For example, undetected DNA swaps in family studies of human disease (e.g., unaffected father is swapped with affected son) will prevent a genetic diagnosis and yield misleading candidate variants. Even without sample errors, the sample manifest (e.g., PED file [1]) containing vital information about the relatedness of individuals within a cohort may include sample-naming errors or swaps. In our experience, familial relationships are often transcribed manually from pedigree diagrams drawn from the clinician. These errors can go unnoticed without careful review.

Therefore, a critical aspect of quality control is assuring that each sequenced DNA sample originated from the expected individual. Unfortunately, these sample-level quality control problems are not detected by existing software leveraging raw sequence data (e.g., FastQC [2]) or sequence alignments (e.g., bam.iobio [3], samtools [4]). While tools such as Plink [1] and KING [5] can detect sex and pedigree errors, they require conversion from the standard VCF [6] format to PLINK format and solely produce text output that requires further manual inspection of custom scripts to detect sample issues. Other tools [7,8] are able to infer pedigree structure from sample genotype data, but they are cumbersome for identifying and resolving sample swaps. To address the need for robust, rapid, and automated detection of problems with sample DNA fidelity, we have developed *peddy*, a software package that evaluates correspondence between the stated sexes, relationships, and ancestries in a pedigree file [1] and those inferred from the genotypes in the VCF file resulting from human WES or WGS studies. *Peddy* is fast and user-friendly: a single command executes a variety of sample analyses directly on a VCF and an associated PED file. The resulting interactive web page and comma-separated (CSV) files describe the results of each quality control test for each sample, as well as indications of which samples are likely to be the result of a mix-up or DNA quality concern.

## Methods

### Overview of quality control measurements

*Peddy* interrogates the genotypes reported in a VCF file to identify potential sample quality problems based on four primary statistics. First, each sample’s stated sex is compared to the genotypes observed on the X chromosome. Second, it compares the degree of relatedness observed between each pair of samples to the expected relatedness measure based on what is stated in the PED file. Third, sample quality is assessed by the count, sequencing depth and ratio of sequence alignments for each allele at sites where an individual is heterozygous. The variance in these measurements facilitates the detection of DNA contamination, unexpected diversity, and insufficient sequencing depth. Finally, the ancestry of the each sample it predicted using a support vector machine (SVM) trained on individuals of diverse ancestry from the 1000 Genomes Project.

### Sampling selected polymorphic sites to increase speed

Each of these quality control statistics can be computationally expensive, especially for whole-genome studies, as standard methods examine every polymorphic locus observed in a study (tens of millions for WGS). Relatedness statistics are especially onerous as they typically require the comparison of each sample to all other samples at each polymorphic site. In order to mitigate the computational cost and improve estimates of relatedness, we have identified a subset of biallelic single-nucleotide polymorphisms (SNPs) from the 1000 Genomes Project (1GP) that mitigate the computational burden incurred in calculating these tests while maintaining high accuracy (Figure 1). Our goal was to identify a subset of markers that are informative for measuring relatedness and predicting ancestry across diverse ancestries and for both whole-genome and whole-exome studies. With these goals in mind, the subset of markers chosen were required to have reported allele frequency greater than 0.04 in each of European, African, American, and South-East Asian populations, a lack of deviation from Hardy-Weinberg Equilibrium (HWE; p-value exceeding > 0.04), and a called (i.e., not unknown) genotype for at least 2,500 of the 2,504 samples in the 1GP. Finally, we required that each locus is also reported to be in the exome capture region common to all of the platforms used in the 1000 Genomes Project so that the set of chosen markers is informative for sample quality measurements in both whole-exome and whole-genome studies. These criteria resulted in a final set of 23,556 biallelic SNPs which are the basis of each statistical measure that *peddy* conducts. In order to quickly interrogate this subset of markers from whole-exome and whole genome studies, *Peddy* performs a tabix [9] genome interval query to each of the sites using cyvcf2 (github.com/brentp/cyvcf2), a python wrapper of htslib (github.com/samtools/htslib) we have developed for programmatic exploration and processing of VCF files.

**Figure 1.**
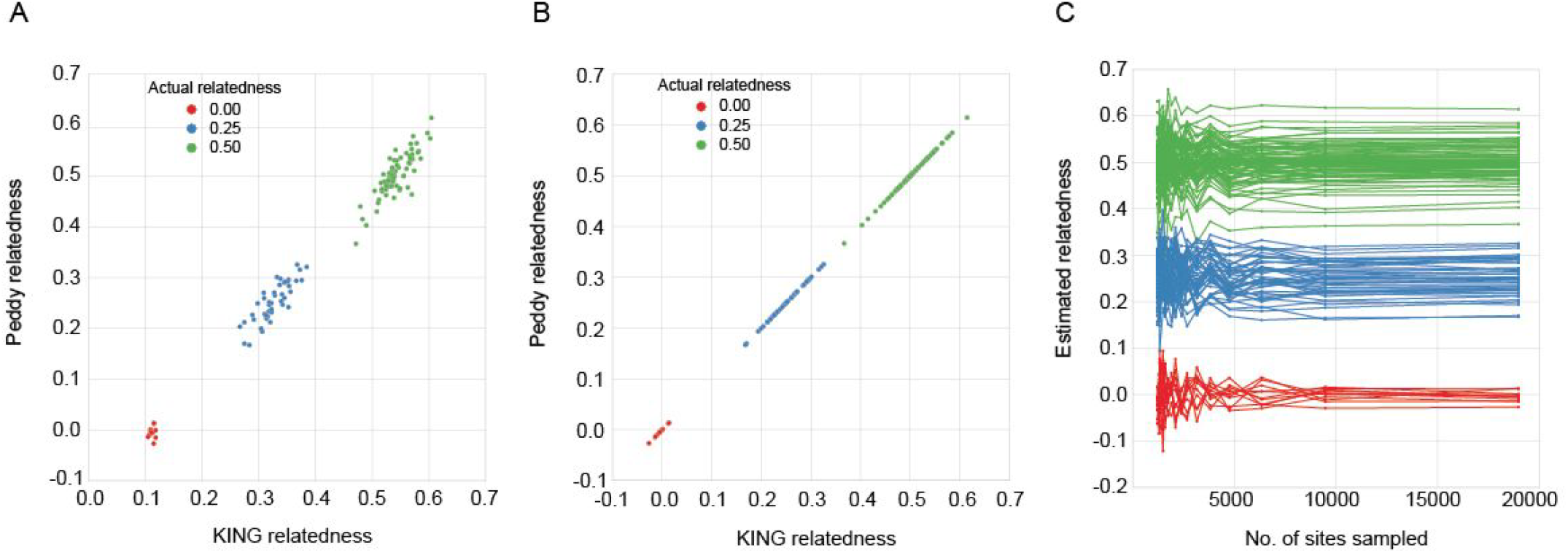
Validation and convergence of sampling method. A comparison of the relatedness coefficient estimated by KING (KING estimates kinship which is 0.5 * relatedness) compared to that from *peddy* **(A)**. A similar comparison when the relatedness estimate is restricted to the subset of 23,556 sites used by *peddy* **(B)**. Convergence of *peddy’s* relatedness estimate as a function of the number of sites sampled **(C)**. The three clusters of converging lines reflect the estimated relatedness among pairs of individuals with an actual relatedness of 0.0, 0.25, and 0.5, respectively. The estimated relatedness rapidly stabilizes to the actual relatedness statistic when at least 5,000 markers are used.

In order to validate the accuracy of our sampling approach, we used the lllumina Platinum Genomes sequencing for the 17 member CEPH pedigree 1463 (ref: [10]) to compare *peddy’s* coefficient of relatedness statistic to the kinship estimate of KING which uses the full set of variants observed. *Peddy*’s coefficient of relatedness is well correlated with KING’s kinship coefficient, and *peddy* provides a better calibrated estimate for unrelated samples from the same family, where we expect the coefficient of relatedness to be approximately 0 (Figure 1A). In our experience, KING generally gives an accurate estimate of kinship, but in this case, it overestimates the relatedness especially for unrelated samples; it has been shown [11] that kinship estimates are more accurate when only sites with less variability in allele frequency among populations are used. When KING is restricted to our subset of 23,556 sites, KING’s estimates are better calibrated (Figure 1B). Furthermore, *peddy*’s estimate of relatedness rapidly converges as the number of sites sampled increases to our full set of 23,556 sites, demonstrating the accuracy and predictive power of the subset of markers we have chosen (Figure 1C). For example, after sampling 10,000 sites, the related estimate converges, implying that ~23,000the full set is a conservatively large. However this larger subset is necessary to maintain accuracy across datasets of diverse quality, as the number of informative sites could be reduced owing to low coverage, quality or exome capture failures. We emphasize that the user may also specify their own selection of sites, thereby making *peddy* usable for any genome and build and allowing it to be applied to studies based upon genotyping arrays or customized research scenarios.

### Pre-processing steps

In addition to the quality control metrics performed by peddy that compare attributes inferred from the genotypes to what is reported in a pedigree file, peddy also performs several internal consistency checks directly on the pedigree file. For example, the pedigree file may report an individual as the maternal parent of another individual, but may also report this individual to have either unknown or male sex. Furthermore, cases often arise in which individuals are listed as parents, yet the parental identifiers are not present as records in the pedigree file. *Peddy* automatically reports these and similar inconsistencies to the user. When provided with a relevant VCF file, *peddy* also reports samples that are present in the VCF, but not in the PED and vice-versa.

### Measures of relatedness

Peddy calculates the coefficient of relatedness from the genotypes observed for each pair of samples using the method described by Manichaikul et al [5] and implemented in the KING software package. We have modified the KING algorithm to use the geometric mean instead of using different formulas for sample pairs from the same family versus those from different families. We have chosen this modification because large pedigrees often contain many unrelated individuals (i.e., via marriage); our results are less affected by this since we’ve chosen sites that are in HWE but we also find that our results more closely match the expected relatedness (e.g., Figure 1A). Our modified formula for the coefficient of relatedness is:

~~~
(N-shared-hets −2 * N-IBSĞ) / 0.5 * geometric_mean(N-hets(i), N-hets(j))
~~~

where *i* and *j* represent the indices for each individual. The algorithmic change with respect to KING’s estimator is that instead of dividing by their sum or the twice the lower number (for the robust estimator), we divide by half of their geometric mean. Otherwise, our relatedness metric is identical to KING’s robust method. The relatedness calculation tests each sample against each other sample so it has an inherent O(n^2^) complexity, but it the computational cost is mitigated by only examining the subset of sites previously described. Furthermore, while majority of the *peddy* codebase is written in Python, this relatedness calculation is written in C for optimal performance. Lastly, we parallelize the computation up to as many processes as are requested by the user. The speed improvement relative to the number of cores scales well, especially as the number of samples increases.

As with KING, our relatedness estimate depends upon the “IBS0” statistic which represents the number of sites at which a pair of individuals shares 0 alleles (e.g., individual *i* is A/A and individual *j* is G/G). The IBS0 statistic is particularly informative when differentiating between parent-offspring and sibling-sibling pairs, as each relationship has an expected coefficient of relatedness of 0.5. In contrast, the IBS0 statistic should be near 0 for parent-offspring pairs since this should only happen in cases of a Mendelian violation (e.g., a *de novo* germline mutation), whereas there should be many such sites observed between siblings (e.g., in cases where both parents are heterozygous and the siblings inherit different alleles from each parent). We also report an “IBS2” statistic, which reflects the number of sites at which both individuals share the same genotype and thus share two alleles. In practice, plotting IBS0 vs IBS2 gives the best visual separation of unrelated samples and grouping of related samples. Nonetheless, we also provide plots of relatedness, since it is a more intuitive metric.

After calculating these statistics, we perform multiple sanity checks based upon the relationships reported in the pedigree file. We ensure that a pair reported as parent-child in the pedigree file has an IBS0 statistic close to zero and alert the user to situations when a pair of individuals has a low IBS0 but is not listed as a parent-offspring pair. Lastly, we report cases in which two individuals are either more or less related than what is stated in the pedigree file. Pairs of samples with a rate of IBS0 less than our empirically derived cutoff of 0.012 are called as parent-child pairs by their genotypes; if they are not indicated as such in the PED file, an error is reported. The information in these reports is also apparent in the interactive plot where points are colored by the relationship defined in the PED file and position according to the inferred relatedness. A blue point (e.g. indicating an unrelated pair) inside a cluster of green triangles (indicating sibling-sibling pairs) is evidence of a sample-swap involving at least one of those samples (e.g., Figure 2C).

**Figure 2.**
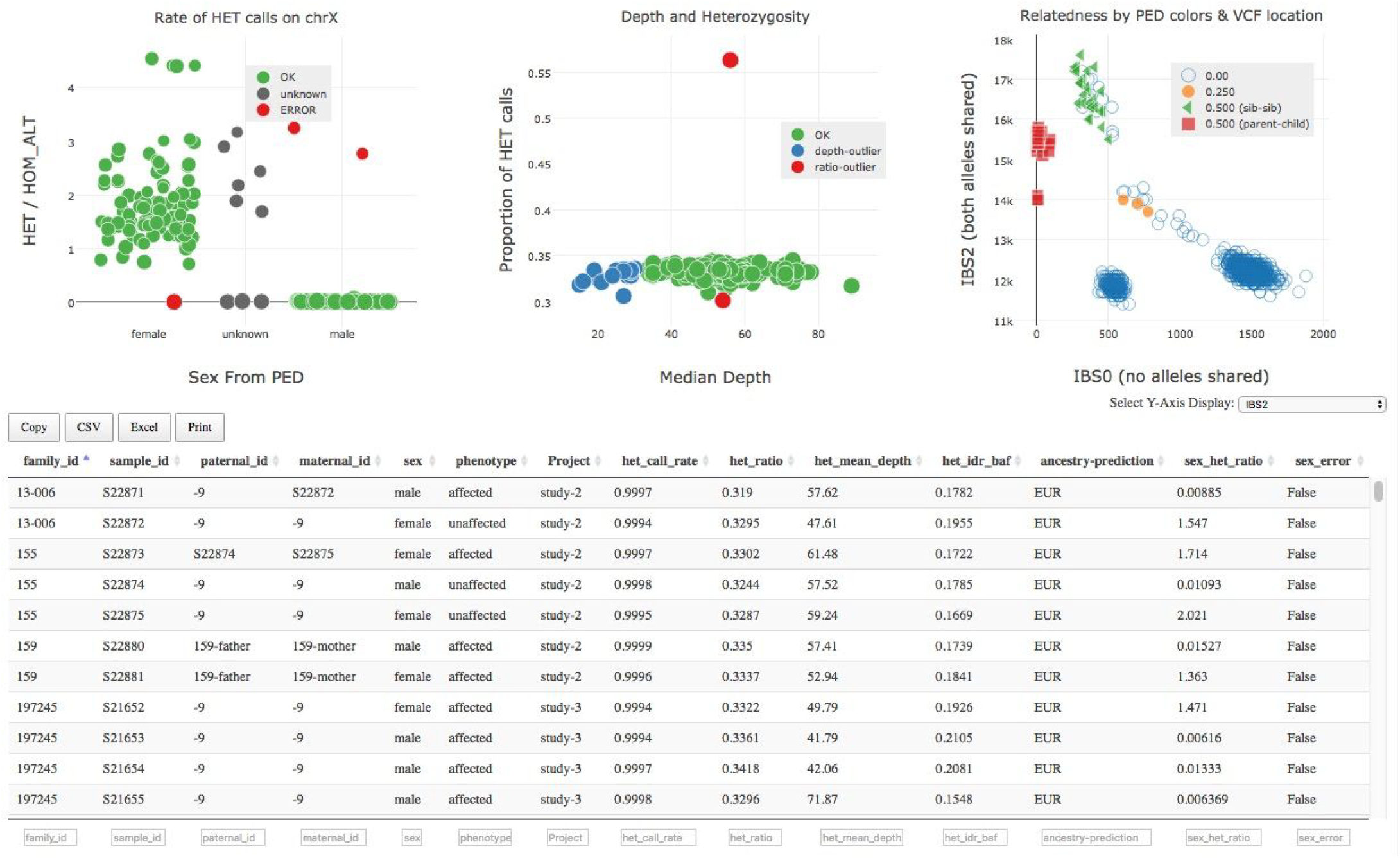
Interactive website for identifying and resolving sample mixups and quality issues. The sex check **(A)**, heterozygosity **(B)**, relatedness **(C)**, and ancestry (not shown; see Figure 4B) plots are interlinked such that clicking in a single point in one plot will highlight all points germane to the selected individual in the other plots. Moreover, the sample information table **(D)** can be sorted, filtered and selected to focus the visualization and interpretation to desired subsets of individuals or families.

### Ancestry prediction

The observed ancestry composition of a cohort is a valuable metric to identify unexpected individuals or potential mix-ups in genetic studies.To predict ancestry, *peddy* leverages the known ancestry of 2,504 individuals collected from diverse world populations as part of the 1000 Genomes Project (a 10 megabyte file of genotypes of these 2,504 individuals is distributed with *peddy*). For each analysis, *peddy* retrains an SVM on the first 4 principal components identified from the encoded genotype vectors (0,1,2,3 for homozygous reference, heterozygous, homozygous alternate, and unknown respectively) of the 1000 Genomes samples using their known ancestries as training labels. This re-training is required since we only include sites that are present above a certain allele frequency and call rate (i.e., the fraction of individuals predicted genotypes at a given site) with among the individuals in a given study. The first four principal components are identified using a randomzed principal component analysis [12] with scikit-learn (see: http://scikit-learn.org/stable/about.html) as a means of dimensionality-reduction. The randomized PCA runs on the 2,504 samples in approximately 4 seconds, making the cost of the re-training negligible.

We decided on 4 principal components and an SVM penalty (C) of 2 using 20-fold cross-validation on the actual 1000 Genomes data with 70 and 30% of samples used to train and test on each split. Those parameters are now used each time to classify incoming samples. Once the SVM is trained on the 1000 genomes samples, the resulting classifier can be applied to the individuals in each study. A study cohort’s individuals are first projected onto the principal components identified from the 1000 Genomes samples and then the SVM is used to predict the ancestry. This information is reported in the text output and in an interactive plot, where the 1000 Genomes samples cluster by “super-population” [13] and the predicted ancestry of the individuals in the cohort is displayed on top of the 1000 Genomes background samples (for an example, refer to Figure 5B). Samples with an SVM prediction probability for a particular ancestry greater than 0.65 are assigned to that ancestry while the remaining samples are classified as unknown.This entire process takes only a few seconds once the genotypes for each site have been collected.

### Sex prediction

By tracking the genotypes observed for each individual outside of the pseudo-autosomal region of the X chromosome, we are able to derive an accurate prediction of an individual’s sex. Since males have only one X chromosome, they should have zero true heterozygote calls in the X chromosome, while females should have a mixture of heterozygous and homozygous genotypes similar to that found in autosomes. We have found that the most informative measure is the ratio of heterozygous to homozygous genotypes. Comparing this ratio to the sex stated in the pedigree file provides another statistic to detect either mix-ups or sex labeling errors in the pedigree file. In cases where the ratio has an intermediate value between the values observed for males and females, it can also be used as an indicator of lower coverage (e.g., a female with a lower heterozygote to homozygote ratio) or individuals with rare sex chromosome disorders such as Klinefelter syndrome [14].

### Detection of poor sequencing quality, consanguinity, or contamination by inspecting the properties of heterozygous genotypes

Examining the heterozygous genotypes and the underlying sequence alignments observed for each individual at each of the 23,556 sampled sites allows *peddy* to detect problems with DNA quality, purity and unexpectedly high levels of homozygosity/heterozygosity. Samples with high rates of heterozygosity or a higher inter-decile range of alternate allele ratios (see Supplemental Figure 1) may be contaminated. Samples with low rates of heterozygosity (and reasonable coverage) could be checked for consanguinity. The data files and visualizations that are generated based upon heterozygous genotypes also report mean observed depth for each sample since the number of heterozygous genotype calls will increase with sequencing depth owing to increased detection power. Finally, *peddy* also reports the per-individual “call rate”, which reflects the proportion of sampled sites with a known genotype, as an indication of the overall quality of an individual’s genotype predictions.

### User interaction

*Peddy* provides text based reports of the above metrics for each individual enabling automated detection of problematic individuals via simple scripts. It also provides an interactive web page (Figure 2) allowing the user to identify potential problems by clicking on points in each plot to inspect which individuals are outliers for each statistic. Each plot is linked so that when an individual point is selected in one plot, the same individual is highlighted in the other plots while also displaying the relevant values of each statistic for that individual. Furthermore, the relatedness plot displays all pairwise measurements comparing the selected individual to other individuals are highlighted. Lastly, the web page provides an interactive HTML table allowing one to filter and sort the displayed individuals by all existing columns provided in the input pedigree file and by the statistics computed by *peddy*. Together, these interactive features provide the researcher with a powerful and efficient means of identifying problematic individuals, possible mix ups, and correcting potential errors in the input pedigree file.

## Results

### Overview

Given a VCF file and associated PED file describing the expected relationships and sex of the individuals in a sequencing study, *peddy* automatically conducts all of the tests described using the subset of 23,556 informative SNPs, thereby allowing the rapid detection of possible issues with individual samples.

To demonstrate *peddy’s* utility for detecting sample issues, we will first demonstrate a contrived example from a small pedigree, followed by a real example derived from a large, whole-genome sequencing cohort collected as part of an ongoing study at the University of Utah. In each case, we will identify discrepancies between what is reported in the pedigree file and what is inferred from the genotypes. The greatest power comes from displaying and integrating all of sample quality control checks together so that one can, for example, note that an individual who appears to be less related than expected to her parents actually has a higher rate of heterozygosity, indicating poor sample quality possibly owing to contamination (which increases heterozygosity and would decrease apparent relatedness). For this reason, we present the results using *peddy's* interactive web-page.

### An intentionally injected error in the CEPH1463 pedigree file

To introduce the functionality in *peddy*, we injected an error into the pedigree file describing a four-member quartet that is a subset of the CEPH1463 17 member pedigree. We intentionally forced a sample swap between the mother (NA12877) and her son (NA12880) by switching the identifiers in the PED file, which is analogous to a situation where the DNA samples were swapped during sequencing. Inspecting the plot resulting from the sex check, we note that one individual stated to be a male (NA12880) and one stated to be a female (NA12877) appear to have the opposite sex indicated by their genotypes (Figure 3A). We can further dissect the potential problem by inspecting the relatedness plot generated by *peddy* (Figure 3B). Because we have four samples, each sample will be involved in a pairing with the three other samples.

**Figure 3.**
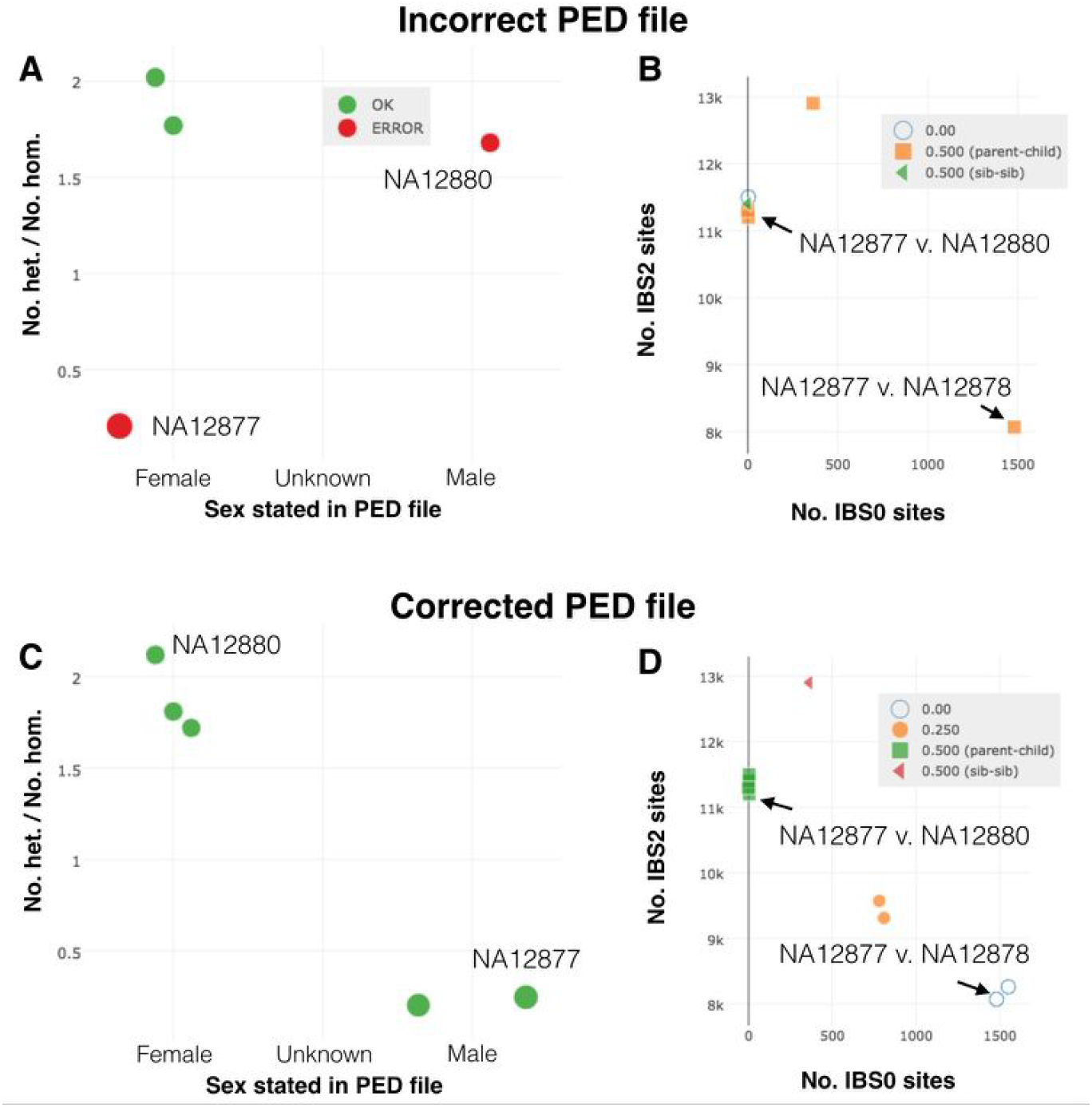
Using peddy to visualize manufactured error in CEPH pedigree. Panel **A** highlights two individuals (red) where the sex stated in the ped file does not match that inferred from the rate of heterozygous calls on the non-pseudo autosomal region of the X chromosome. In the relatedness plot **(B)**, we can see that the swap has caused unexpected relationships (or lack thereof) for both individuals. In panels **C** and **D** we can see that these errors have been resolved by switching the names of the sample in the PED.

This means that the two sample errors we have created propagated as errors in the observed relatedness with all other individuals. The simplest error to follow is one where a the pedigree file indicates a parent-child relationship, yet the genotypes do not support such a relationship. For example, the number of IBS0 sites observed between individual NA12877 and NA12878 is much higher than expected for a true parent-child relationship (that is, near 0). Since Figure 3A indicates a sex swap, we can further infer from the relatedness information that NA12877 and NA12880 had been swapped. When we rerun *peddy* with a corrected pedigree file reflecting this knowledge, we observe the expected sex and relatedness (Figure 3C,D). In our experience, such examples are even simpler to resolve as most studies deal with smaller families such as trios and quartets and the interactive web page allows us to filter to specific families. Since each *peddy* run completes in seconds, these samples errors can be fixed iteratively by retesting with *peddy* after each correction in order to resolve all potential errors stepwise fashion.

### Unexpected heterozygosity rates and leveraging ancestry predictions

As part of a large, ongoing study of rare, familial diseases, we have whole-genome sequenced multiple families at the University of Utah. We applied *peddy* to a subset of 225 individuals from this study in effort to further demonstrate the types of errors that it can detect. While we are not able to share the data due to HIPAA constraints, we use this cohort to demonstrate the utility in a large study. First, the plot of heterozygous genotypes reveals a single individual (S15084) with a substantially higher rate of heterozygote calls than any other individual (Figure 4A). While a small increase in the rate of heterozygosity could suggest a different ancestry, in this case, the level seen here is extreme. In addition, we can use the PCA ancestry plot to confirm that individual S15084 is predicted to be of European ancestry and clusters well many other individuals of the same ancestry (Figure 4B). Therefore, we conclude that this DNA sample may be the result of contamination with one or more other samples, and we may therefore remove this sample from downstream analyses. Similarly, individual S21051 is also an outlier exhibiting a lower rate of heterozygosity. Such a lack of heterozygosity could be caused by either extremely low average sequencing coverage (thereby minimizing the power to detect heterozygous genotypes) or consanguinity. Since this plot also reveals that individual S21051’s mean sequencing depth observed across all heterozygous genotypes is 49.1, it is possible that the lack of heterozygosity arises from consanguinity, so we may want to follow up on this individual and family by talking with the researcher or examining the relatedness that *peddy* reports for the parents.

**Figure 4.**
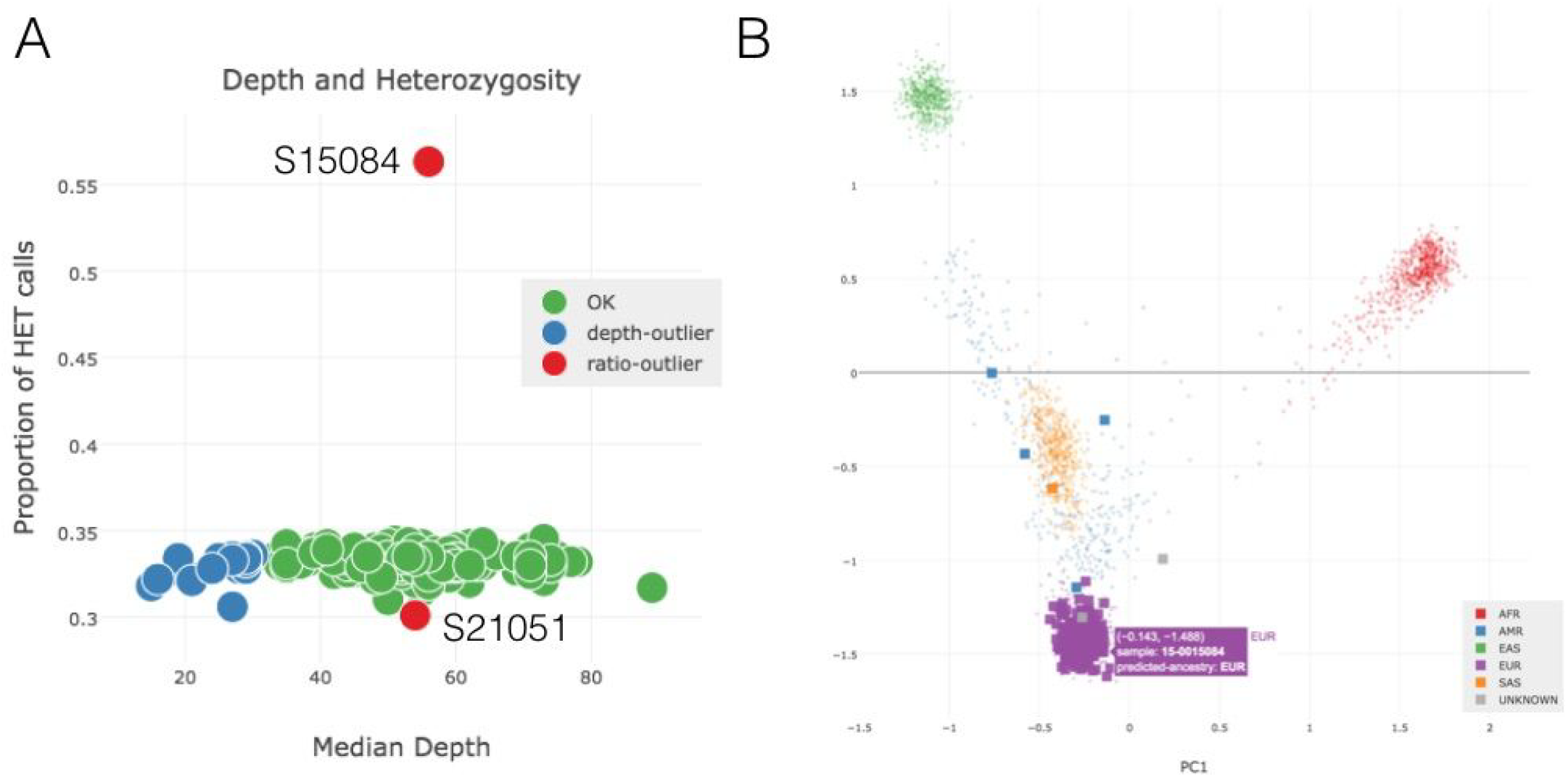
Depth and Heterozygosity and Ancestry. Outlier individuals with unexpectedly high and low proportions of heterozygous (HET) genotypes. **(B)** A PCA analysis is conducted and an SVM trained on the 1000 Genomes samples (small background points) is used to predict the ancestry of each of the individuals in a study (large square points).

### Examples of unexpected measures of relatedness

*Peddy* also produces an interactive plot of different measures of relatedness (i.e., IBS0, IBS2, coefficient of relatedness). This relatedness plot colors each point by the expected relatedness based upon the relationship declared in the pedigree file. The location of each point in the plot is determined by the relatedness measures inferred from the genotype data for each individual in the VCF file. As expected, given that the 225 samples came from many different families, most of the points on the plot reflect pairs of unrelated individuals (blue). However, we observe a large cluster of blue points with a lower-than expected IBS0 (Figure 5A; asterisk). If we hover over those points we see that S15084 is a member of all of those pairs. This observation is consistent with the prior observation that this individual has a much higher number of heterozygous genotypes (Figure 4A). Therefore, this outlier individual shares the alternate allele more frequently with other individuals, thereby reducing the number of IBS0 sites observed with other samples. Individual S15084 is also a member of the red cluster of points with IBS0=0 but below the larger red cluster (Figure 5A; ampersand). This highlights the utility of the interactive plots; a problematic sample will typically appear as an outlier in several of the plots.

**Figure 5.**
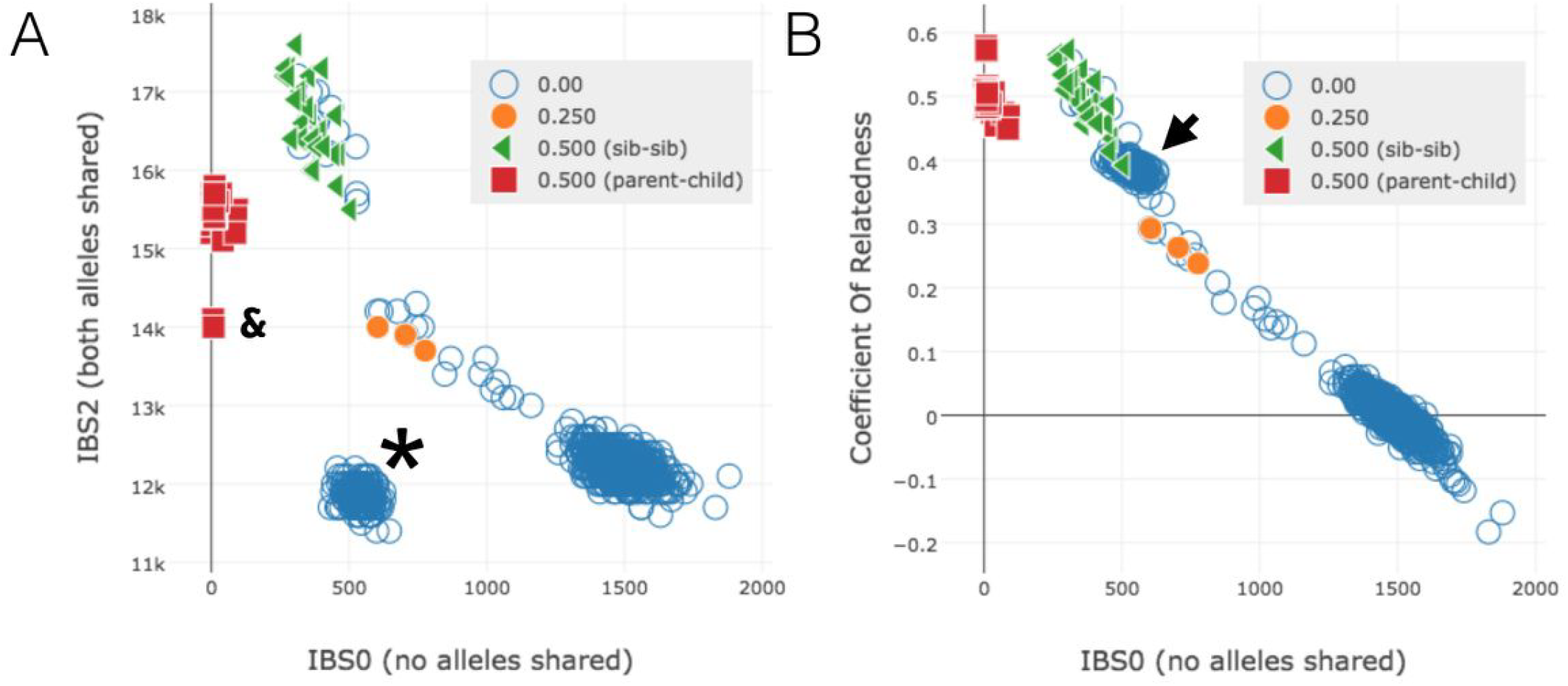
Relatedness with IBS2 or CoR. We compare plots with IBS2 **(A)** or the coefficient of relatedness **(B)** for the same data. The coefficient of relatedness gives and intuitive value to check that, for example, siblings have a CoR of 0.5 and unrelated pairs have a CoR of around 0. But IBS2 often provides better visual separation of clusters even with lower quality data. The cluster of blue points with an IBS0 around 500 and IBS2 around 12K are clearly unrelated in the IBS2 plot **(A)** but in the relatedness plot, they appear to cluster almost with the cluster of sib-sib pairs (green triangles). This blue cluster is all from a single sample with a high rate of heterozygote calls that skews the relatedness calculation.

In the cluster of sibling-sibling pairs indicated by green triangles, we see a number of blue circles, indicating that the pedigree file does not explicitly identify several pairs that are actually siblings. In most of these cases, we go back to the pedigree file and see that the siblings were sequenced and their parents were not added to the pedigree file or were unspecified for those individuals. Similarly in the case of the yellow circles, we find that in nearly all cases, these individuals are stated as belonging to the same family, yet the details of the family structure were not specified in the pedigree file.

The relatedness plot provides the ability to represent the y-axis as either IBS2 (Figure 5A;) or the coefficient of relatedness (Figure 5B). While coefficient of relatedness (CoR) is a more intuitive metric, we find that IBS2 provides greater separation, thereby allowing the researcher to identify samples issues with greater ease. For example, both sibling-sibling and parent-child relationships have the expected CoR (Figure 5B). However, the CoR plot leads us to believe that the cluster of unrelated relationships with individual S15084 exhibits a high degree of relatedness, on par with sibling-sibling pairs (Figure 5B; arrowhead). In contrast, the IBS2 plot clearly illustrates that the relationships with individual S15084 are aberrant (Figure 5A; asterisk). This example motivates the complementary utility of both metrics for evaluating unexpected relatedness.

Finally, we can tie those insights in with what we can ascertain from the sex plot. We see that a number of samples didn’t have their sex specified in the pedigree file and therefore appear in the center as grey (Figure 6A; X-axis, “unknown”). In such cases, *peddy* will assign the sex predicted by the ratio of heterozygous to homozygous genotypes on the X chromosome. We also find an obvious swap between two parents in a trio. This results in one red point for individual S22867 and another for individual S22868, where the genotypes from the X chromosome do not match the sex reported in the pedigree file. By leveraging the interactive sample selection feature, we observe that these individuals each exhibit the expected relatedness to the child; it is only the genotypes observed on the X chromosome that provide the evidence for the clear sample swap.

**Figure 6.**
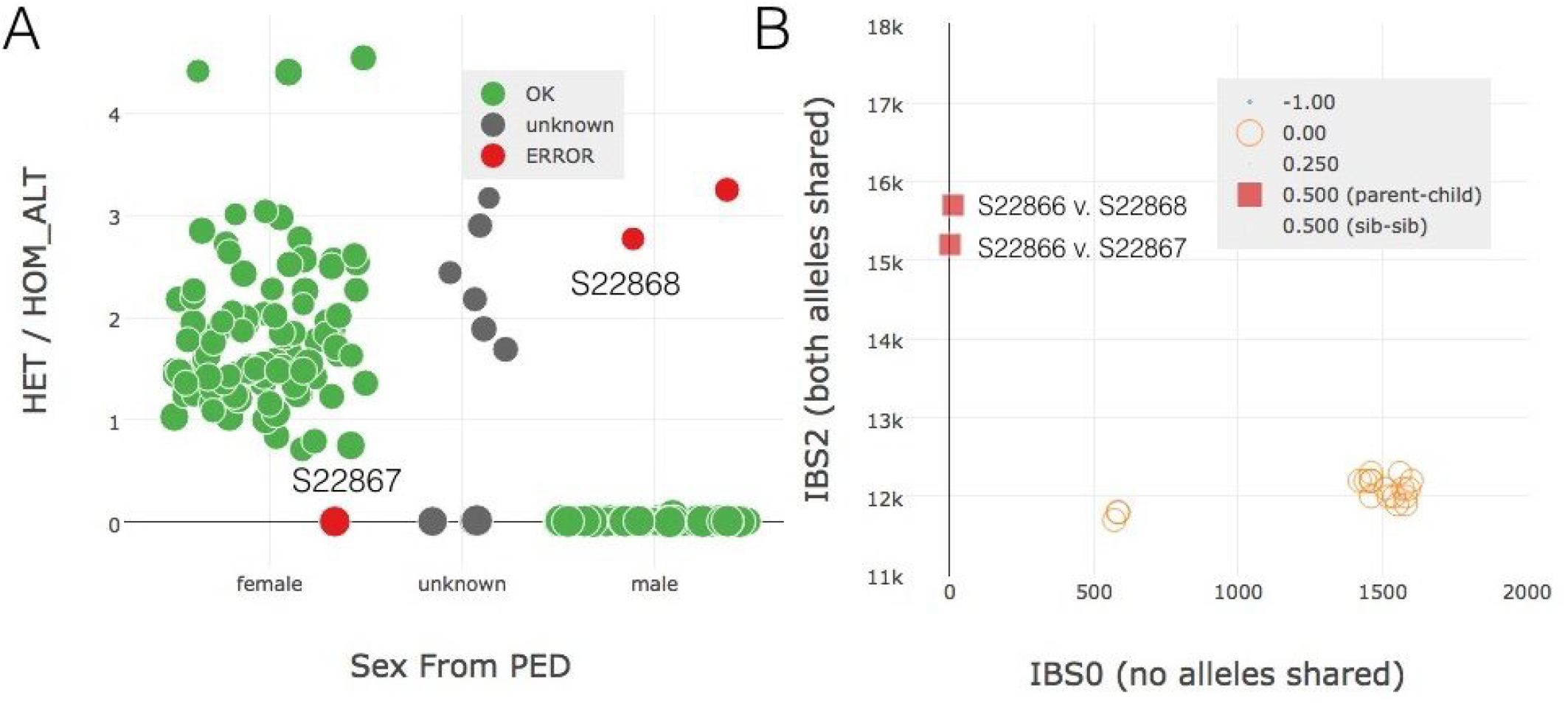
Sex Plot and Sample Selection. Upon seeing an potential sample-swap in panel **A** with two members from the same family, we can leverage the table selection tool (not shown) to highlight solely that family in the relatedness plot **(B)**. In so doing, we verify that this is a husband-wife pair where both have the expected relation to their child. This allows one to infer that the husband and wife labels have been swapped.

## Conclusions

*Peddy* is a powerful software package that facilitates the detection and correction of sample quality issues and mix-ups that complicate analysis and inhibit discovery. Its interactivity substantially improves upon the functionality available in previous software packages and the reports it produces allows for the development of simple scripts to automate sample quality control measures. While not presented here, since *peddy* infers ancestry, relatedness and sample sex, we emphasize that it may also be used to add information to PED files of anonymous DNA samples. Lastly, *peddy’*s efficiency and flexibility allow it to be used for a broad range of studies, ranging from studies of family trios to large-scale investigations of thousands of human genomes. As such, we anticipate that *peddy* will be a vital tool for quality control in current and future human genome and exome studies.

## Acknowledgements

Barry Moore provided valuable feedback on early versions of the software and suggested the idea of predicting an individual’s ancestry. Daniel McGoldrick and Jessica Chong provided helpful feedback and QC on the software.

## Author contributions

BSP conceived of and implemented the software, analyzed the data, and wrote the manuscript. ARQ conceived of the software and wrote the manuscript. All authors read and approved the final manuscript.

## Supplementary Materials

Assuming minimal bias during DNA library preparation, the ratio of sequence alignments harboring the alternate allele is expected to follow a binomial distribution (p∼0.5) for all sites at which an individual is heterozygous (Supp Figure 1A, bottom panel). Substantial deviation from this expectation is potential evidence for either aberrantly low average sequencing depth or contamination with DNA from other individuals in the DNA library (Supp Figure 1A, top panel). *Peddy* measures the interdecile range (10th to 90th percentile; IDR) of alternate allele ratios from heterozygous genotypes as a statistic to summarize the degree to which the binomial expectation is violated for each individual (Supp Figure 1B). Individuals with potential contamination will have substantially more heterozygous genotypes than other individuals and will have a higher alternate allele ratio IDR. In contrast, individuals conceived from consanguineous parents will have substantially fewer heterozygous genotypes, reflecting a higher degree of homozygosity.

**Supplemental Figure 1.**
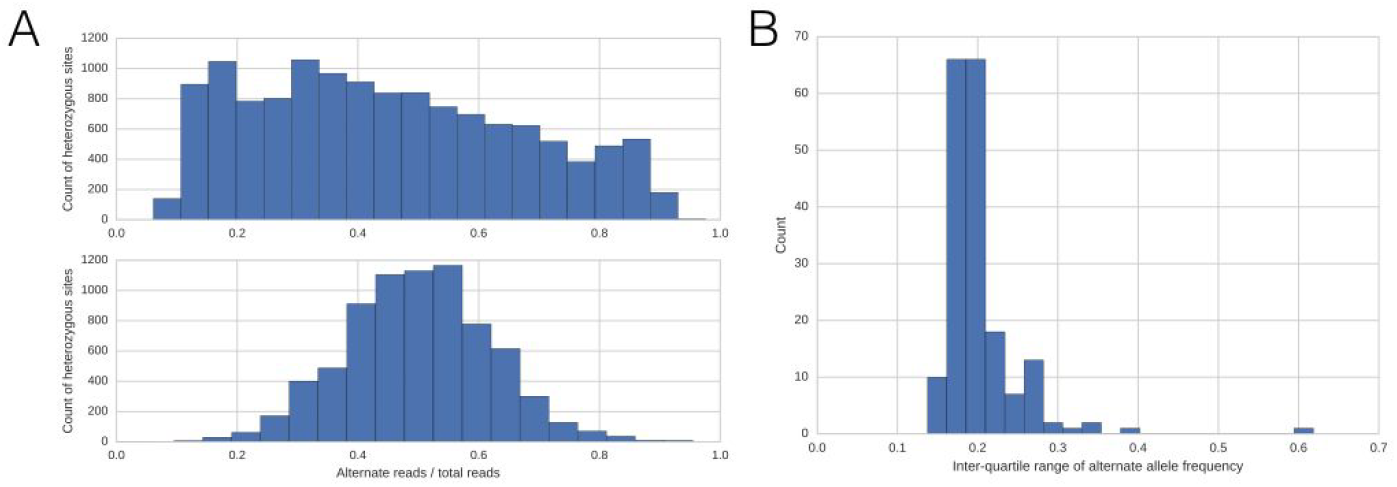
Interdecile range of fraction of alternate reads. The top panel in A shows a sample with a large inter-decile range while the bottom panel shows the distribution of a good-quality sample with a lower range. The distribution of all samples is shown in Figure 1B, where we can clearly see the outlier at the far right.

